# Optimizing an Automated Processing Pipeline for Regional Macromolecular Proton Fraction and Volume Measurements in the Brain

**DOI:** 10.64898/2026.09.21.753351

**Authors:** Neva M. Corrigan, Daniel S. Hippe, Vasily L. Yarnykh

**Affiliations:** Institute for Learning & Brain Sciences, University of Washington, Seattle WA, USA; Fred Hutch Cancer Center, Seattle, WA, USA; Department of Radiology, University of Washington, Seattle WA, USA

**Keywords:** Macromolecular proton fraction, quantitative MRI, myelin imaging, brain parcellation, reproducibility, reliability

## Abstract

There is increasing demand for quantitative measurements in neuroimaging research, including absolute quantification of regional myelin density. Fast whole-brain macromolecular proton fraction (MPF) imaging enables acquisition and reconstruction of quantitative maps of MPF, a biomarker of myelin, and has been used in many research applications. To date, no standard protocols have been published for parcellating MPF maps. FreeSurfer is a widely used software package for automated brain parcellation. It is designed for use with high-resolution T1-weighted anatomical images, typically acquired with an MPRAGE sequence. MPF maps inherently provide high contrast between white and gray matter and can potentially be used as input images for parcellation, enabling simultaneous quantitative tissue characterization and volumetric assessment. However, it is unclear how the use of MPF maps as input images with FreeSurfer software affect reliability and repeatability of regional MPF and volume estimates. We evaluated three FreeSurfer parcellation workflows for this purpose. The workflows differed in the amount of neuroimaging data required and computational intensity. MPF and MPRAGE data were acquired in two separate sessions for 11 adults. Reliability was evaluated using mean relative differences in estimates, Bland-Altman analysis, and intra-class correlation coefficients (ICCs). Repeatability was assessed using within-subject coefficients of variation (CVws). All three workflows produced similarly high within-subject repeatability across scans. However, reliability in regional MPF estimates was lower in gray matter than in white matter for workflows that used MPF maps as input. Workflows that use MPF maps as FreeSurfer input may be adequate for estimating mean MPF in GM and WM of cortical parcels and in subcortical regions. However, the MPRAGE-based workflow is recommended when reliable individual-level estimates or volume estimates are required.

## Introduction

Regional brain myelination is affected by factors such as developmental change, environmental influences, pathophysiology, and aging. Macromolecular proton fraction imaging (MPF) is increasingly being used to capture these changes. Studies in animal models have found to be MPF to be strongly correlated with histological markers of myelin (Khodanovich et al., 2018; Kisel et al., 2022). MPF has been used to characterize patterns of changes in human brain myelination during typical development, including in the fetal brain (Korostyshevskaya et al., 2018, 2019; Yarnykh et al., 2018), the infant brain (Corrigan et al., 2022; Zhao et al., 2022) and the adolescent brain (Corrigan et al., 2021). In infants, regional brain myelination has been found to be predictive of later language development (Corrigan et al., 2022) and executive function (Zhao et al., 2022). MPF has been found to be predictive of symptom severity in bipolar disorder and depression (Gusakova et al., 2026), schizophrenia (Smirnova et al., 2021; Sui et al., 2022), multiple sclerosis (Yarnykh et al., 2015), chronic pain (Filimonova et al., 2025), and long-COVID (Khodanovich et al., 2025).

Calculation of MPF is based on a two-pool magnetization transfer (MT) model that describes magnetic energy exchange between a semi-solid macromolecular and a mobile water proton pool. A key parameter of this model is MPF, which reflects the relative amount of macromolecular protons that contribute to the MT effect. The single-point MPF mapping method (Yarnykh, 2012, 2016) allows for acquisition of whole-brain MPF data in scan times that are practical for clinical and research purposes. In this method, an MT-weighted volume, a T1-weighted volume, and a proton-density-(PD)-weighted volume are acquired. Additional acquisition of B0 and B1 field maps is optional. B0 field correction can be omitted in human brain imaging at clinical field strengths due to a negligible effect of B0 field inhomogeneity on MPF (Yarnykh et al., 2020). Correction of transmit B1 field inhomogeneity, while generally necessary, can be accomplished without additional data acquisition using a recently developed data-driven method (Yarnykh, 2021).

To date, there are few guidelines for optimal post-processing of MPF maps, as well as little previous work that addresses the repeatability of regional mean MPF estimates derived using different registration and parcellation protocols. MPF maps can be analyzed using whole-brain voxel-wise methods, or by averaging across regions derived from manual or automatic parcellation. Automatic parcellation is particularly useful for rapid processing in large studies, and for producing consistent measurements as compared to manual approaches. FreeSurfer is a widely used software for automatic parcellation of T1-weighted images. High-resolution T1-weighted MPRAGE images are commonly used as input for FreeSurfer processing because their high gray-white matter contrast and isotropic 3D acquisition are well suited to surface-based morphometric analysis (FreeSurfer, 2009). It is unclear whether the high gray-white matter contrast and isotropic 3D acquisition of MPF maps themselves might be adequate to be substituted as input images into FreeSurfer for calculation of both regional mean MPF and volume estimates.

In this context, we evaluate three processing workflows for deriving mean regional estimates of MPF and regional volumes, ordered here from highest to lowest processing complexity. The MPRAGE workflow requires a high-resolution MPRAGE T1-weighted volume acquired in the same scan session as the MPF data. FreeSurfer is used to parcellate the MPRAGE volume, and the MPF map is registered to the parcellated volume to calculate regional mean MPF and volume estimates. The MPFreg workflow entails performing registration of the MPF component images prior to MPF map reconstruction and then inputting this map directly into the FreeSurfer software. The MPF workflow omits registration of the component images prior to MPF map reconstruction, and uses the resulting map as input to FreeSurfer. The goal for this study was to evaluate the effect of these three different automated workflows on the reproducibility and scan-rescan repeatability of regional mean MPF and volume estimates.

## Methods

### Sample

Data were collected from 11 adults with no known neurological, psychiatric, or major somatic illness (6 females, 5 males; mean age ± SD = 44.1 ± 11.9 years, range 28-65 years). Each participant was scanned at two time points, separated by a mean interval of 296 days (SD = 148 days, range 28 - 447 days). MPF data were acquired for all participants at both time points. MPRAGE data were acquired at both time points for six participants and at only one time point for the remaining 5 participants. Informed consent was obtained from all study participants. All study procedures were approved by the Institutional Review Board at the University of Washington, and informed consent was obtained from all study participants.

### Data acquisition

All data were acquired on a 3.0 T Philips Achieva MRI system using an 8-channel head coil. High-resolution T1-weighted images of the head were acquired using an MPRAGE sequence with FOV = 240 × 240 × 180 mm, acquisition voxel size 1.0 × 1.0 × 1.0 mm^3^, reconstructed voxel size 0.5 × 0.5 × 0.5 mm^3^, TR/TI/TE = 5.9/900/2.8 ms, shot interval 2500ms, flip angle (FA) = 8°, and scan time 4 min 45 s. A 3D MPF mapping protocol was implemented according to the single-point synthetic reference method (Yarnykh, 2012, 2016) and included three spoiled gradient-echo sequences with MT (TR = 28ms, flip angle (FA) = 10°), proton-density (TR = 21ms, FA = 4°), and T1 (TR = 21ms, FA = 25°) contrast weightings. Off-resonance saturation in the MT-weighted sequence was applied at the offset frequency 4 kHz with effective FA = 560° and pulse duration 12 ms. The images were obtained in the sagittal plane with dual-echo readout (TE1/TE2 = 2.3/6.9 ms), FOV = 240 × 240 × 180 mm^3^, and actual voxel size of 1.25 × 1.25 × 1.24 mm^3^ interpolated to 0.625 × 0.625 × 0.620 mm^3^. Parallel imaging (SENSE) was used in two phase encoding directions with acceleration factors of 1.5 and 1.2. In all sequences, non-selective excitation and optimal spoiling with the excitation pulse phase increment of 169° (Yarnykh, 2010) were used. The total scan time for acquisition of MPF mapping data was about 15 min.

### MPF map reconstruction

MPF maps were reconstructed according to a single-point synthetic reference algorithm (Yarnykh, 2016) with automatic data-driven correction of B1 field non-uniformity (Yarnykh, 2021) using custom-written C-language software with previously determined (Yarnykh, 2012) constraints for the non-adjustable two-pool model parameters. Surrogate B1 field reconstruction was performed with previously published algorithm constants r_0_ = 0.3 and r_f_ = 4.5 (Yarnykh, 2021). Correction of B0 field inhomogeneity was not applied because of a negligible effect of B0-related errors on MPF measurements (Yarnykh et al., 2020). Prior to map reconstruction, individual echo images in each data set were averaged to increase SNR (Helms & Dechent, 2009). The MPF maps are comprised of voxel-wise estimates of the macromolecular proton fraction, expressed as a percentage of the total proton content.

For each participant and scan, MPF maps were reconstructed using two separate workflows, hereafter referred to as the MPF and MPFreg workflows. In the MPFreg workflow, image volumes were registered using rigid-body registration with the FLIRT toolbox of the FSL software package (v. 6.0.5; Smith et al., 2004) prior to MPF map reconstruction. In the MPF workflow, no registration of the component image volumes was performed prior to MPF map reconstruction.

MPF maps from both workflows were reoriented to match the orientation of the corresponding scanner-reconstructed MPRAGE images. Images from the MPRAGE volume acquired for one participant and scan session are shown in Figure 1A. Images from MPF map created with the MPFreg workflow for the same participant and scan session are shown in Figure 1B.

**Figure 1.**
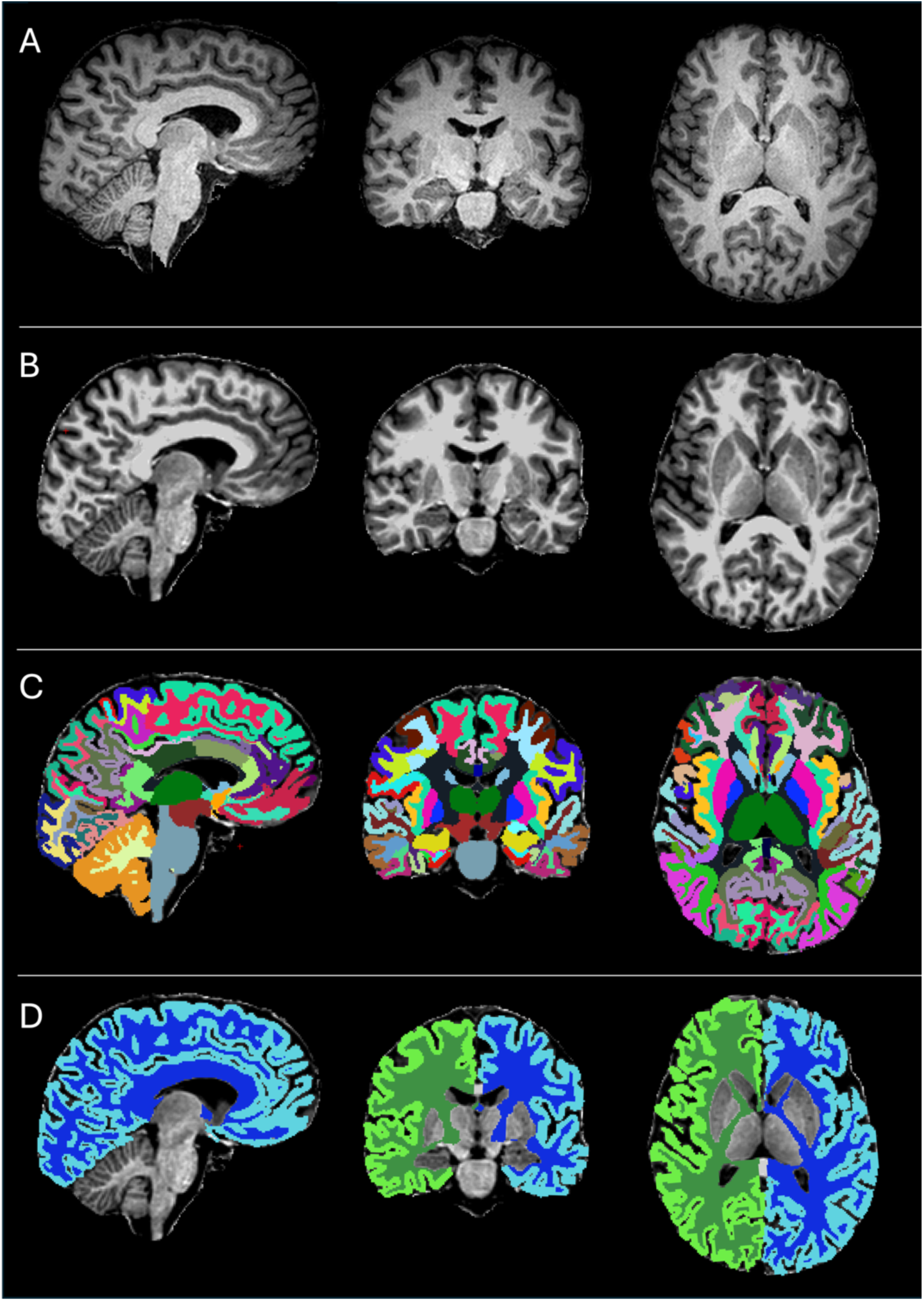
Example orthogonal slices from the MPRAGE volume, MPF map, parcellation results, and cerebrum masks from the same scan session for a single participant. A. MPRAGE volume. B. MPF map calculated using the MPFreg workflow. C. GM and WM parcels and subcortical region masks. D. Cerebrum GM and WM masks. Note that parcels and masks were derived from the MPF map in B.

### Automated parcellation

The Desikan-Killiany (DK) Atlas (Desikan et al., 2006) was selected as the parcellation template within FreeSurfer. The DKT atlas provides 34 cortical gray matter (GM) parcels and 34 corresponding cortical white matter (WM) parcels in each hemisphere. FreeSurfer also provides segmentation of subcortical structures. Subcortical regions included in the present analysis were the bilateral nucleus accumbens, amygdala, caudate, cerebellum cortex, cerebellum white matter, hippocampus, pallidum, putamen, thalamus, ventral diencephalon, the brainstem, and 5 corpus callosum segments (anterior, central, mid-anterior, mid-posterior, posterior).

For comparison of the effect of different inputs to FreeSurfer, parcellation was performed using three workflows: MPF, MPFreg and MPRAGE. The MPFreg and MPF workflows used the corresponding MPF map as the FreeSurfer input volume. The MPRAGE workflow used the MPRAGE volume as input. The “recon-all” function of the FreeSurfer software package (v.8.0; Fischl, 2012) was used to parcellate the input data. Since the MPF map reconstruction software performs skull stripping, for the MPFreg and MPF workflows, the “-noskullstrip” argument was used to skip the FreeSurfer skull stripping step. An example parcellation produced by FreeSurfer for a single participant using the MPFreg workflow is shown in Figure 1C.

### Construction of masks

Masks for left and right frontal, temporal, parietal and occipital lobes were created by combining the parcels associated with each lobe from the wmparc.mgz file produced by FreeSurfer. Separate lobe masks were made for GM and WM. The frontal lobe masks were constructed by combining the superior frontal, rostral and caudal middle frontal, pars opercularis, pars triangularis, pars orbitalis, lateral and medial orbitofrontal, precentral, paracentral, frontal pole, rostral anterior cingulate, and caudal anterior cingulate parcels. The parietal lobe masks were constructed by combining the superior parietal, inferior parietal, supramarginal, post-central, precuneus, posterior cingulate, and isthmus cingulate parcels. The temporal lobe masks were constructed by combining the superior, middle, and inferior temporal parcels, as well as the banks of the superior temporal sulcus, fusiform, transverse temporal, entorhinal, temporal pole, and parahippocampal parcels. The occipital lobe masks were constructed by combining the lateral occipital, lingual, cuneus, and pericalcarine parcels. Hemispheric and whole-brain GM and WM masks were similarly created. For each hemisphere, a GM mask was constructed by combining the 34 GM parcels regions in the ribbon.mgz file output by FreeSurfer for that hemisphere. A WM mask for each hemisphere was created by combining the 34 WM parcels from wmparc.mgz with the regions labeled “UnsegmentedWhiteMatter”. Example GM and WM masks are shown in Figure 1D. A schematic of FreeSurfer input images, and calculations of parcellations and masks for each of the three workflows is shown in Figure 2A.

**Figure 2.**
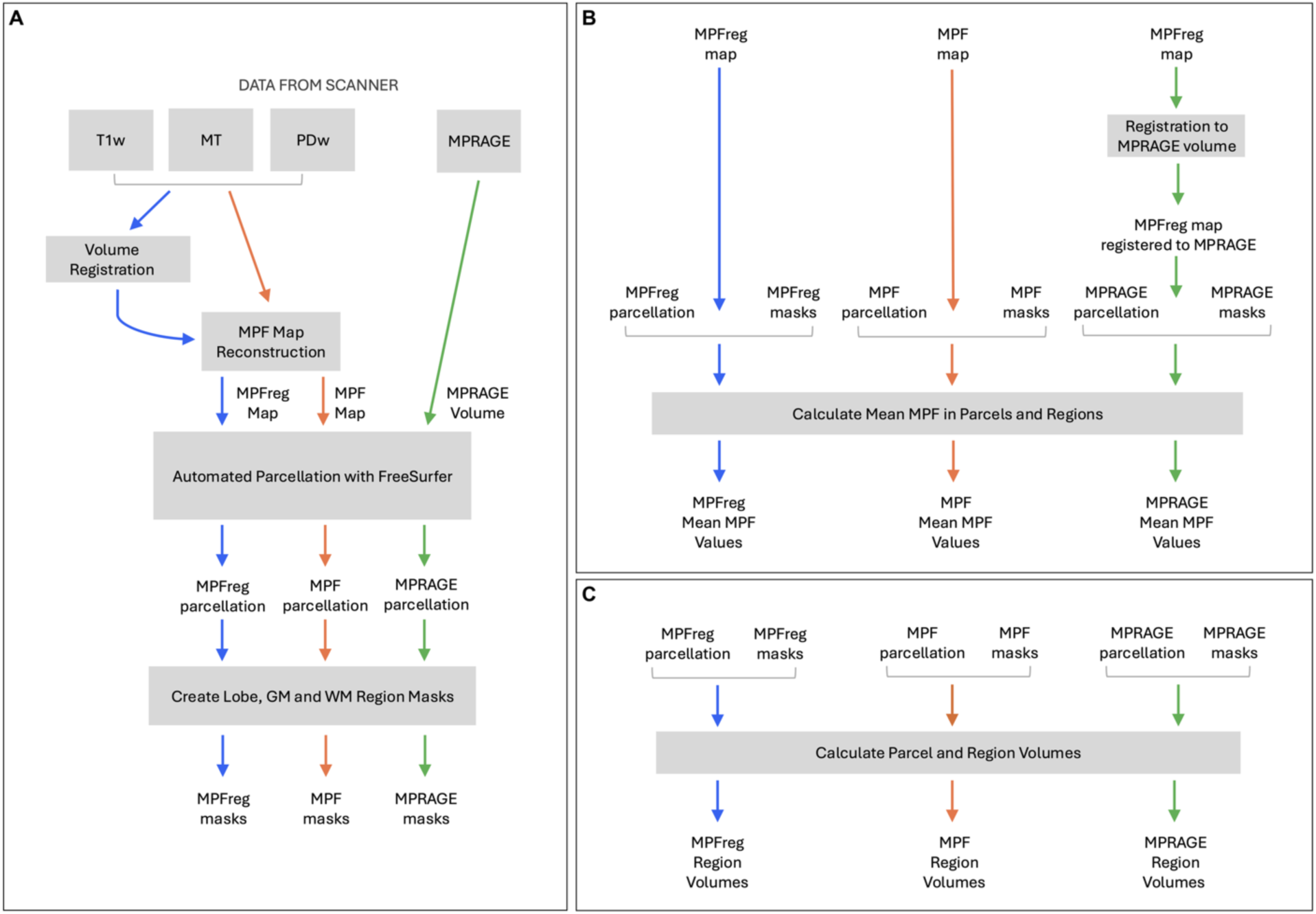
Workflows for (A) automated parcellation and generation of regional masks, (B) calculation of mean MPF values within parcels and regions, and (C) calculation of parcel and region volumes. In all panels, blue represents the MPFreg workflow, orange represents the MPF workflow, and green represents the MPRAGE workflow.

### Calculation of mean regional MPF

For the MPRAGE workflow, prior to calculation of mean myelin density, the MPF maps from the MPFreg workflow were registered to the “brain.mgz” volumes output by the FreeSurfer processing of the MPRAGE data using the Advanced Normalization Tools (ANTs) software package (v.2.6.2; Avants et al., 2011). For all workflows, the FreeSurfer “mri_segstats” function was used to calculate mean MPF values. For cortical GM and WM parcels, the wmparc.mgz file was specified as the parcellation volume, and for lobe and hemisphere regions, the region mask was specified as the parcellation volume. To ensure inclusion of only voxels in GM, WM or subcortical structures, all voxels with MPF values below 2% were excluded from the calculation of mean MPF. A schematic for calculation of mean regional MPF values for each workflow is shown in Figure 2B.

### Calculation of regional volumes

For each workflow, the “aparcstats2table” function of the FreeSurfer software package was utilized to extract the volume for each cortical GM parcel from the FreeSurfer processing output. The “asegstts2table” function was used to extract the volumes of subcortical regions from the FreeSurfer processing output. Volumes were additionally calculated using this function for the GM and WM hemisphere regions. Volumes were not calculated for cortical WM parcels or WM lobes because the boundaries of WM parcels are based on proximity to cortical GM parcels rather than independently defined by WM anatomy. A schematic for calculation of regional volumes for each of the three workflows is shown in Figure 2C.

### Statistical analysis

We calculated and compared reliability and repeatability of results generated by the three workflows. We use the term reliability to refer to agreement in measurements obtained using different processing workflows applied to data from the same participant in the same scan session. We use the term repeatability to refer to the consistency of measurements obtained using the same processing workflow across repeated acquisitions from the same participant (test–retest).

The primary reliability metrics we used were the mean difference (bias), the limits of agreement (LOA) and intraclass correlation coefficient (ICC). The LOA were calculated as the mean difference ± q x (standard deviation of the difference), where q is the 97.5% percentile of the t-distribution with degrees of freedom = n – 1 (Bland & Altman, 1986), and was summarized as the width of the interval (wLOA). Mean difference and LOA were calculated on absolute difference and relative difference scales. Relative difference was calculated as the difference in natural logs (i.e., ln[X_1_] - ln[X_2_]) as a symmetric approximation to the percentage difference. The ICC was calculated using a two-way mixed-effects model using the *irr* package in R (version 0.84.1; Gamer et al., 2026). The consistency ICC, which summarizes the within-subject variability relative to the between-subject variability with bias excluded, was used. This metric does not provide overlapping information with the mean difference summary of bias. We used the following guidelines for interpreting ICC values: poor (0-0.49); moderate (0.5-0.74), good (0.75-0.9), excellent (>0..9; Koo & Li, 2016).

Reliability metrics were calculated separately for each region, hemisphere, and tissue compartment. To simplify presentation and interpretation, the metrics from left and right hemispheres for each region were averaged. Regional metrics were displayed and ranked using forest plots. Aggregate GM, WM, and subcortical region Bland-Altman plots were used to display the mean differences of all regions within each category. All reliability metrics were calculated using the 17 scans with MPRAGE acquisitions from the 11 participants. Each scan of the same participant was treated as a separate but not independent observation. Confidence intervals (CIs) and p-values were calculated using the non-parametric bootstrap with resampling by participant to account for non-independence between scans of the same participant (Huang, 2018). P-values were adjusting for multiple testing across regions using the Benjamini-Hochberg method to control the false discovery rate (FDR; Benjamini & Hochberg, 1995).

The primary scan-rescan repeatability metric was the within-subject coefficient of variation (CVw), which is a measure of the consistency, or precision, of repeated measurements from the same participant on the same imaging system within the same scan session (van Houdt et al., 2024). The CVw was expressed as a percent and calculated using the root mean square method (Chubb et al., 2018; Hyslop & White, 2009). As with reliability, repeatability metrics were calculated separately for each region, hemisphere, and tissue compartment, with the metrics for the left and right hemispheres averaged. Repeatability metrics for all three workflows were calculated using the 6 participants with two scans with MPRAGE acquisitions. CIs and p-values were calculated using the non-parametric bootstrap with resampling by participant. P-values were adjusting for multiple testing using the Benjamini-Hochberg method, separately from the reliability analysis.

Statistical calculations were performed using R version 4.2 (R Foundation for Statistical Computing, Vienna, Austria). All CIs and hypothesis tests were two-sided with statistical significance defined as adjusted p < 0.05. All CIs are 95% CIs without adjustment for the number of comparisons.

## Results

### Agreement between MPF measurements for processing workflows

#### Mean MPF

##### Regional mean MPF values

Mean relative difference in mean regional MPF values between workflows for the cerebrum, lobes, cortical parcels, and subcortical regions in shown in Figure 3. In WM, the MPF workflow showed significantly larger mean MPF values than the MPFreg workflow in the cerebrum, all 4 lobes, and all 34 cortical parcels. Both the MPF and MPFreg workflows showed significantly larger mean MPF values than the MPRAGE workflow in all WM regions and parcels.

**Figure 3.**
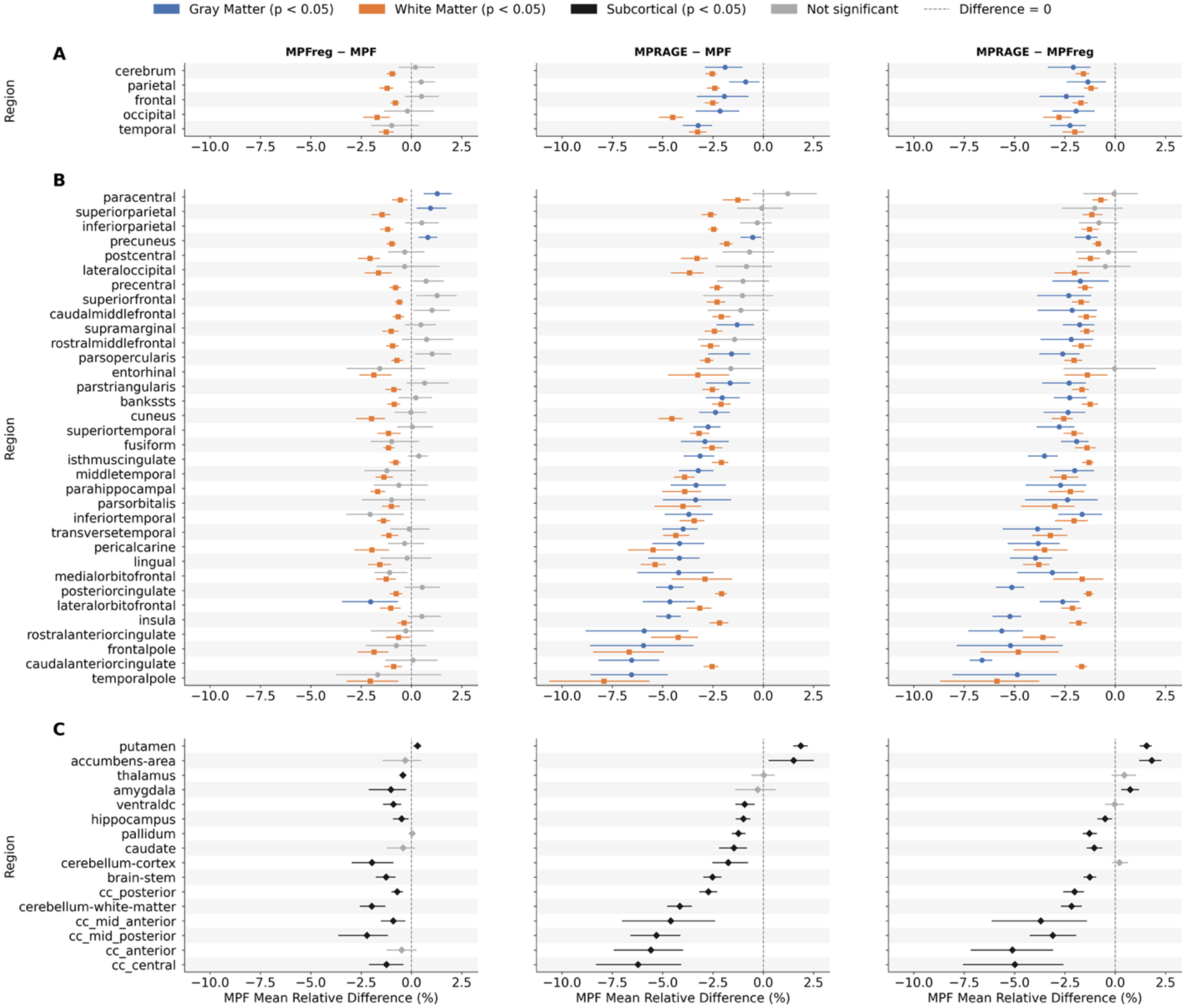
Mean relative difference in mean MPF values between processing workflows for GM and WM in cerebrum and lobe regions (A), cortical parcels (B), and for subcortical regions (C). Markers indicate mean difference and horizontal lines represent 95% confidence intervals. Colored and black markers indicate statistically significant differences. Gray markers indicate non-significant differences.

In GM, no significant differences in mean MPF values between the MPF and MPFreg workflows were found in the cerebrum and lobe regions. In the cortical parcels, significant differences were found for four cortical parcels (with three parcels showing larger MPFreg than MPF values and one parcel showing a larger MPFreg than MPF value). Both the MPF and MPFreg workflows showed significantly larger mean MPF values than the MPRAGE workflow in the cerebrum and all lobes, and in a subset of cortical parcels (24 for MPF and 28 for MPFreg).

Mean MPF values were significantly larger for the MPF workflow than for the MPFreg workflow for 11 of the 16 subcortical regions. One subcortical region, the putamen, had significantly higher mean MPF values for the MPFreg than the MPF workflow. Mean MPF was larger for the MPF workflow than the MPRAGE workflow in all subcortical regions except the thalamus and amygdala, which showed no significant differences, and the putamen and nucleus accumbens, which showed larger relative values for the MPRAGE workflow. Mean MPF was larger for the MPFreg workflow than the MPRAGE workflow in 10 of the 16 subcortical regions. The putamen, nucleus accumbens, and amygdala showed larger mean MPF values for the MPRAGE workflow than for the MPFreg workflow, and the thalamus, ventral diencephalon and the cerebellum cortex showed no significant differences.

##### Intra-Class Correlation Coefficient (ICC)

ICC values for workflow comparisons for mean MPF values across brain regions are shown in Figure 4. ICC values for the comparison between MPF and MPFreg workflows were greater than 0.90 in all WM cerebrum, lobe and parcel regions. ICC values for comparison of the MPFreg and MPF workflows to the MPRAGE workflow were greater than 0.90 for all WM cerebrum and lobe regions, with the exception of the occipital lobe for the MPF v MPRAGE comparison (ICC=0.89). ICC values were generally high across cortical parcels for both comparisons, with most exceeding 0.90 and nearly all exceeding 0.75. ICC values in the WM for comparisons to the MPRAGE workflow were lowest for the frontal and temporal pole parcels.

**Figure 4.**
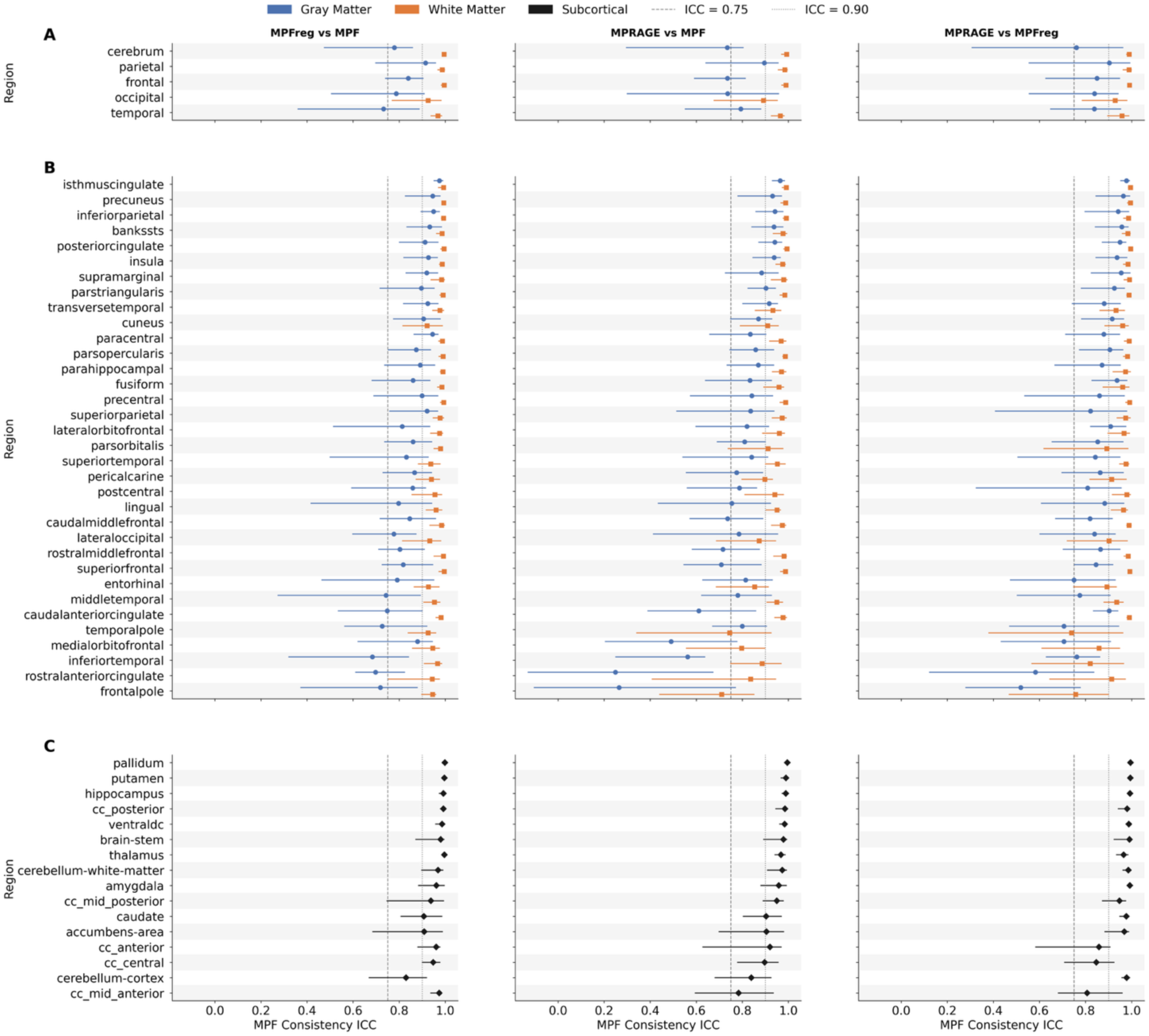
ICC for mean regional MPF values between processing workflows for GM and WM in cerebrum and lobe regions (A), cortical parcels (B), and for subcortical regions (C). Markers show average ICC and horizontal lines represent 95% confidence intervals. Vertical lines indicate ICC thresholds of 0.75 (dashed line) and 0.90 (dotted line), conventionally interpreted as good and excellent reliability, respectively.

ICC values for all workflow comparisons were smaller in GM regions than for corresponding WM regions. GM ICC values exceeded 0.75 for 29 of the 34 cortical parcels for the MPF vs. MPFreg comparison, 26 of the 34 cortical parcels for the MPF vs. MPRAGE comparison, and 29 of the 34 cortical parcels for the MPFreg vs MPRAGE comparison. For all comparisons, regions of the brain with the lower ICC values tended to be those near air-tissue boundaries. These included the frontal pole, rostral anterior cingulate, inferior temporal, temporal pole, and medial orbitofrontal cortex.

Subcortical ICC values for all method comparisons exceeded 0.75. Subcortical ICC values exceeded 0.90 in 15/16 regions for the MPF vs MPFreg comparison. For the MPF vs. MPRAGE comparison, subcortical ICC values exceeded 0.90 in 13/16 regions, and for the MPFreg vs. MPRAGE comparison, they exceeded 0.90 in 13/16 regions.

##### Bland-Altman Analysis

Bland-Altman plots for absolute differences in mean MPF for comparisons between workflows are shown in Figure 5. The analysis revealed a consistent ordering of mean MPF estimates, with the MPF workflow yielding the highest values, the MPRAGE workflow yielding the lowest values, and the MPFreg workflow yielding intermediate values. The magnitude and variability of these differences depended on tissue type. Mean bias between the MPF and MPFreg workflows was greater in cortical WM than in GM and subcortical regions. Comparisons of these workflows to the MPRAGE workflow showed both greater negative bias and wider wLOA, with the highest wLOA in subcortical regions. Average biases were small compared to the widths of the limits of agreement (wLOA), suggesting that region-to-region variability is greater than the systematic average bias. When expressed relative to the mean MPF value across workflows for each tissue type, the wLOA ranged from 11% to 15% in cortical GM, from 5% to 9% in cortical WM, and from 6 to 12% in subcortical regions, with the highest relative agreement observed for the comparison of the MPF workflow to the MPFreg workflow.

**Figure 5.**
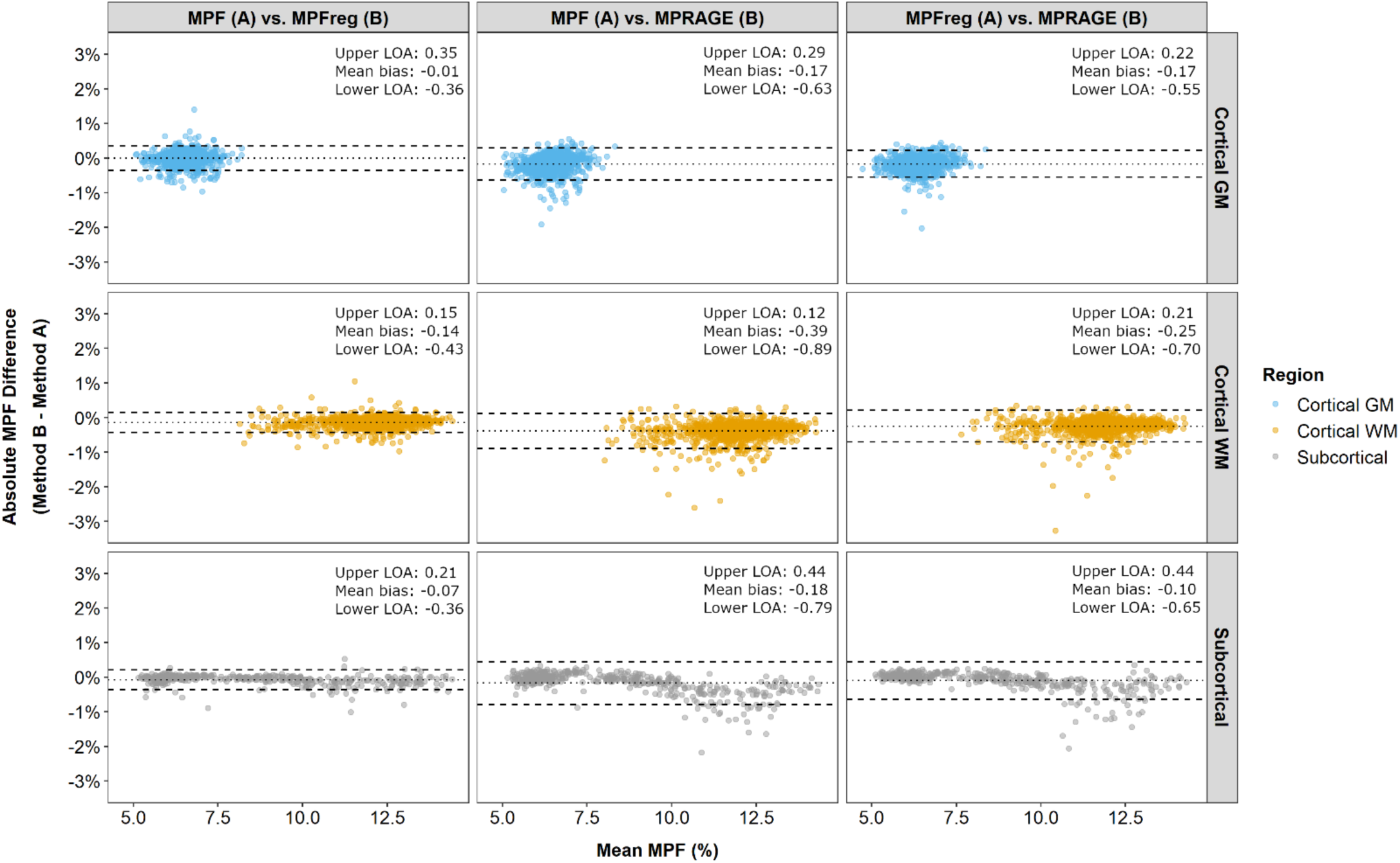
Bland-Altman plots for absolute difference in mean MPF comparisons between workflows for cortical GM and WM parcels, and for subcortical regions. Dotted and dashed lines show mean bias and 95% limits of agreement (LOA), respectively.

Comparisons of absolute MPF wLOA between workflows for the GM and WM of the cerebrum, lobes, and cortical parcels are shown in Figure 6. For the comparison between MPF and MPFreg workflows, GM wLOA were larger than those for WM in the cerebrum, 3 of the 4 lobes, and 30 of the 34 parcels. For the comparisons of the MPF and the MPRAGE workflow, wLOA was greater in the GM than the WM in the cerebrum, 3 of the 4 lobes and 23 of the 34 parcels, and for the MPFreg comparison to MPRAGE, wLOA was greater for GM than WM in the cerebrum, 3 of the 4 lobes, and 22 of the 34 parcels.

**Figure 6.**
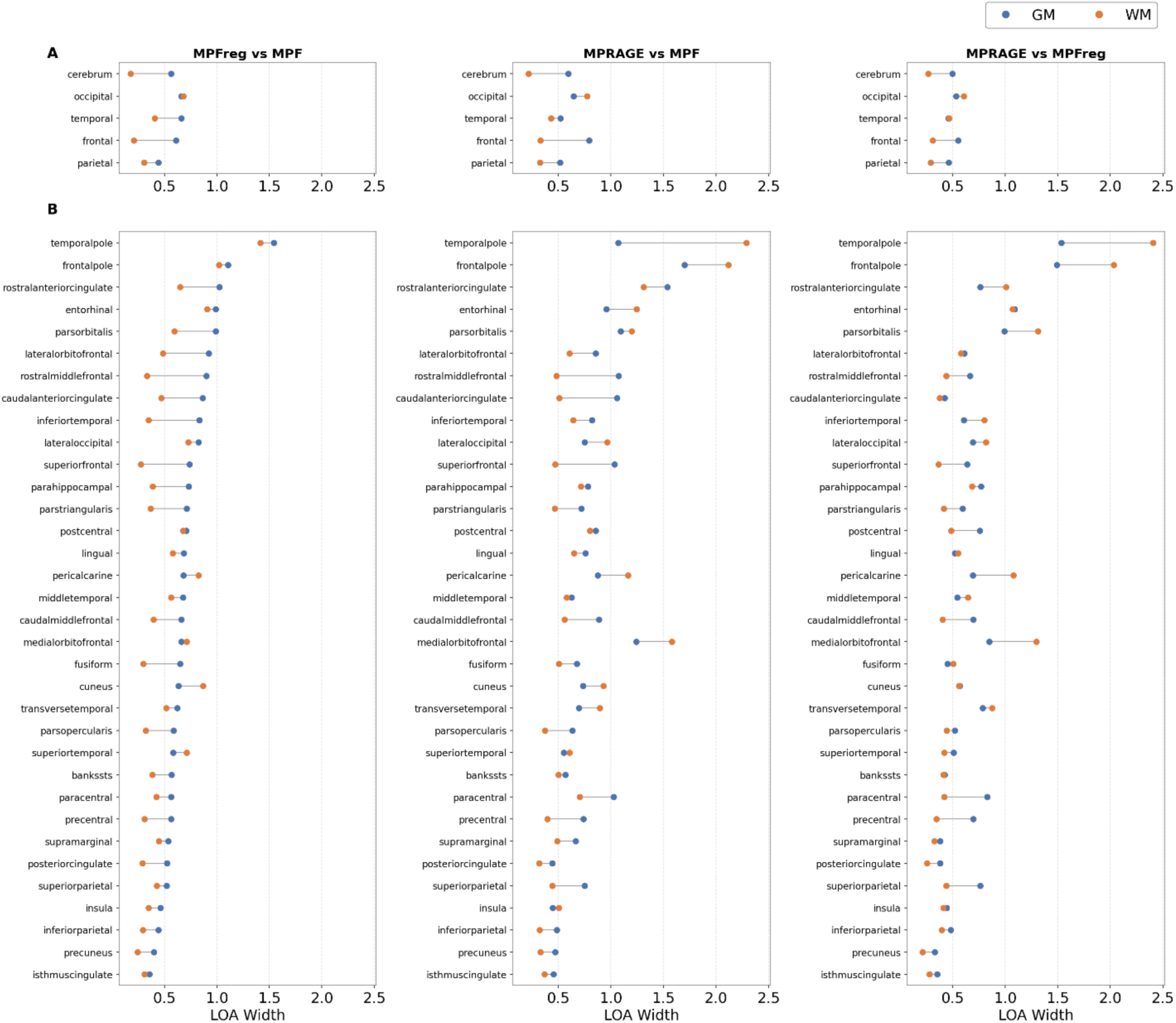
Plots of absolute widths of the limits of agreement (wLOA) in mean MPF from Bland-Altman analysis of workflows for GM and WM in the cerebrum and lobe regions (A) and cortical parcels (B).

#### Volumes

##### Regional volume values

Relative volume differences between the workflows for the cerebrum, lobes, cortical parcels, and subcortical regions are shown in Figure 7. For volume calculation, the only WM region evaluated was the cerebrum. The MPRAGE workflow resulted in cerebrum WM volumes that were higher than for both the MPF and MPFreg workflows. MPFreg workflow cerebrum volumes were larger than those of the MPF workflow.

**Figure 7.**
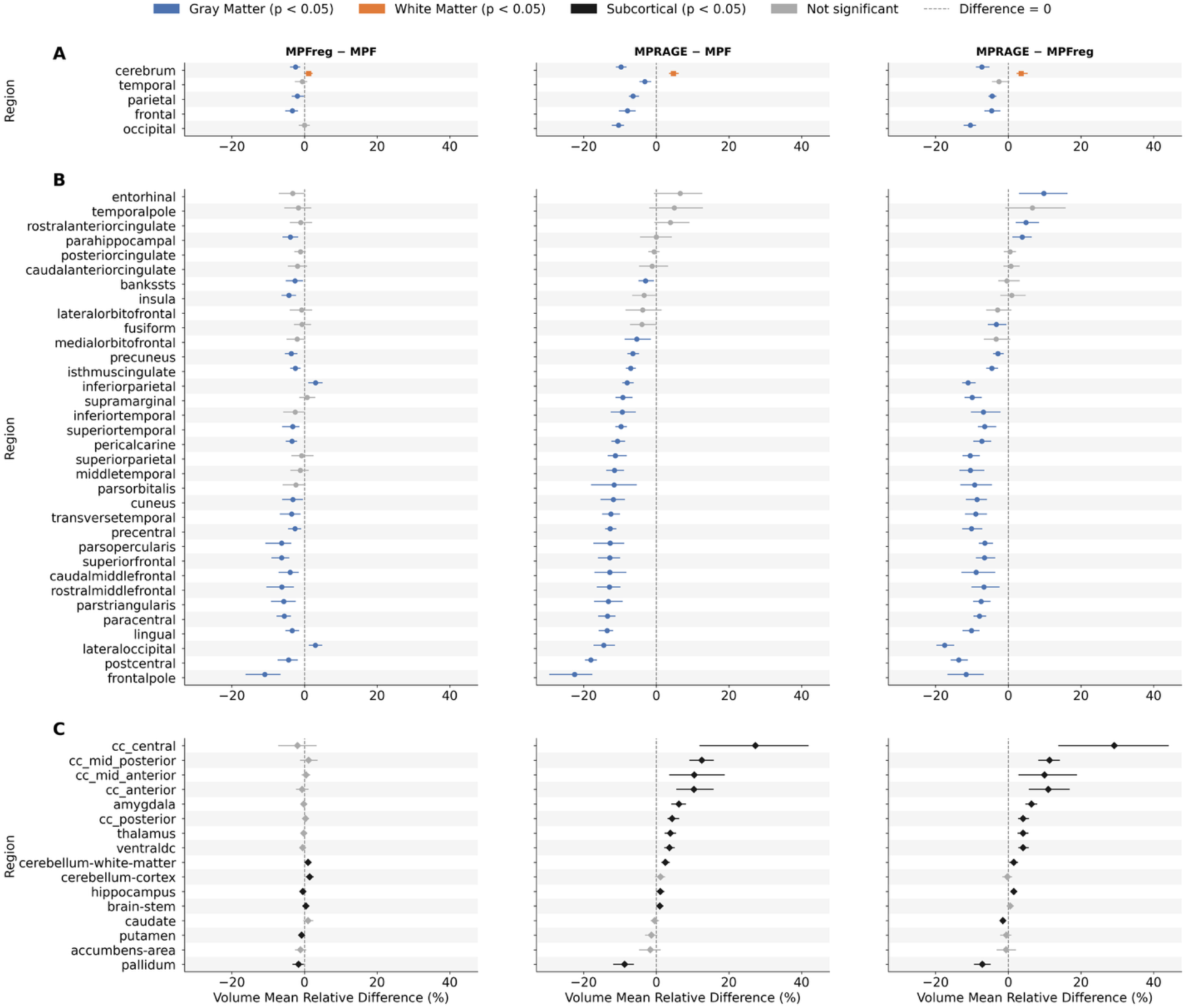
Mean relative difference in volume between processing workflows for the cerebrum and lobe regions (A), cortical parcels (B), and subcortical regions (C). Markers indicate mean difference and horizontal lines represent 95% confidence intervals. Colored and black markers indicate statistically significant differences. Gray markers indicate non-significant differences.

In GM, the MPF workflow resulted in significantly larger volumes than the MPFreg workflow in the cerebrum, 2 of the 4 lobes, and 19 of the 34 cortical parcels. The MPF workflow resulted in significantly smaller volumes than the MPFreg workflow in 2 cortical parcels. Both the MPF and MPFreg workflows showed significantly larger mean volumes than the MPRAGE workflow in the majority of cortical parcels. For the MPFreg vs. MPRAGE comparison, volumes were significantly larger for the MPRAGE workflow for 3 cortical parcels.

For the subcortical regions, the MPFreg workflow resulted in significantly different volumes than the MPF workflow in 6 of the 16 subcortical regions, with smaller volumes in 3 regions and larger volumes in the other 3. Mean volumes were larger for the MPRAGE workflow than the MPF workflow in 11 subcortical regions and for the MPFreg workflow in 10 subcortical regions. In contrast, volumes for the pallidum were smaller for the MPRAGE workflow than for both the MPF and MPFreg workflows.

##### Intra-Class Correlation Coefficient (ICC)

ICC values for relative volume differences are shown in Figure 8. The ICC values for the cerebrum WM were greater than 0.90. In GM, ICC values were greater than or equal to 0.90 in the cerebrum and all lobes, with the exception of the parietal lobe in the MPF vs. MPFreg comparison (ICC = 0.89) and the temporal lobe for the MPFreg vs. MPRAGE comparison (ICC = 0.88). GM ICC values exceeded 0.75 for 32 of the 34 cortical parcels for the MPF vs. MPFreg comparison, 23 of the 34 cortical parcels for the MPF vs. MPRAGE comparison, and for 32 of the 34 cortical parcels for the MPFreg vs MPRAGE comparison. The temporal and frontal pole parcels had the lowest ICC values for comparisons to the MPRAGE workflow.

**Figure 8.**
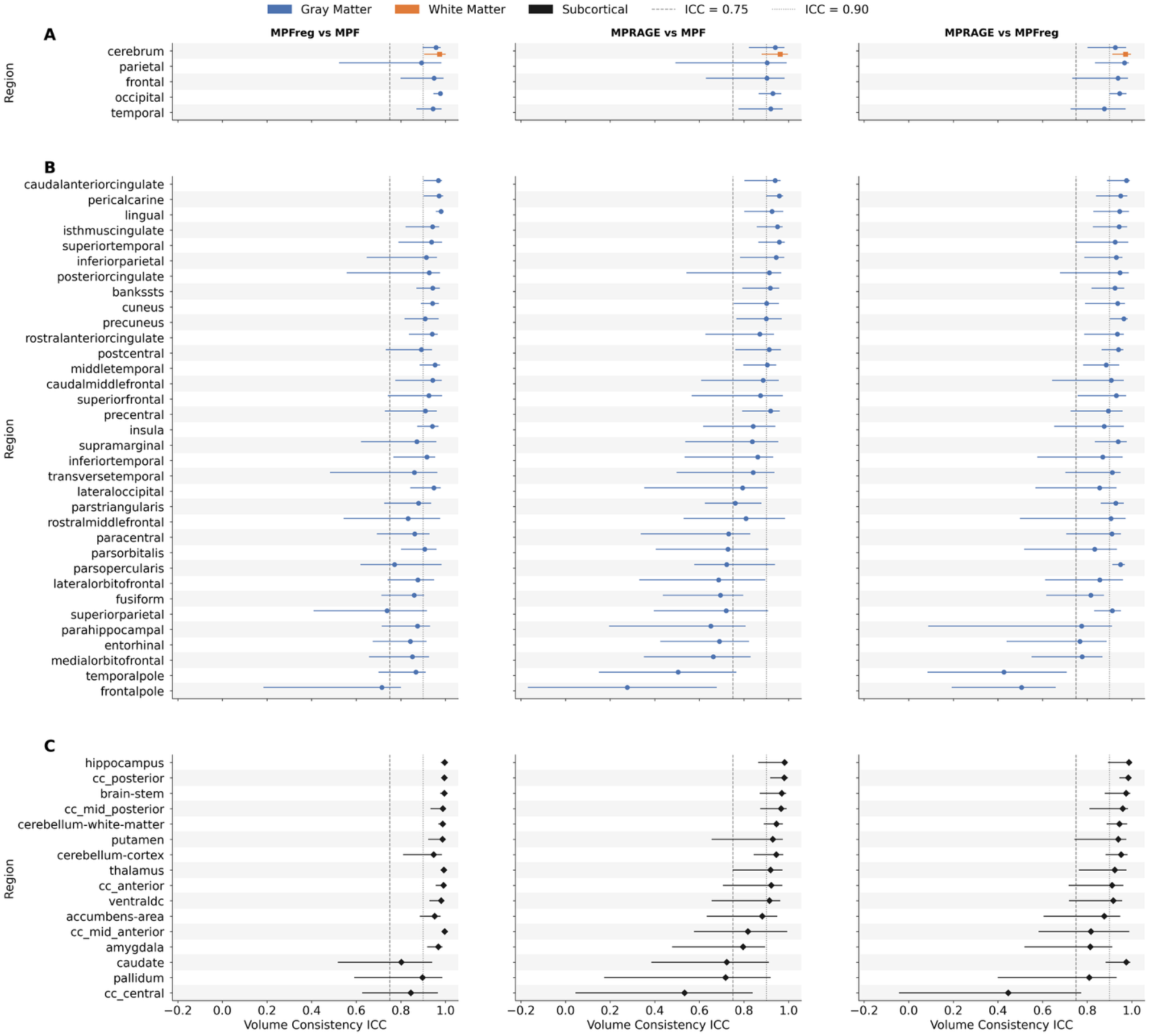
ICC for volumes between processing workflows for cerebrum and lobe regions (A), cortical parcels (B), and subcortical regions (C). Markers show average ICC and horizontal lines indicate 95% confidence intervals. Vertical lines indicate ICC thresholds of 0.75 (dashed line) and 0.90 (dotted line), conventionally interpreted as good and excellent reliability, respectively.

Subcortical ICC values exceeded 0.90 in 13/16 regions for the MPF vs MPFreg comparison. For the MPF vs. MPRAGE comparison, subcortical ICC values exceeded 0.90 in 10/16 regions, and for the MPFreg vs. MPRAGE comparison, they exceeded 0.90 in 11/16 regions.

##### Bland-Altman Analysis

Bland-Altman plots for relative differences in regional volumes between workflows are shown in Figure 9. For cortical GM, the MPF workflow yielded average volumes that were greater than the MPFreg workflow, which in turn yielded average volumes that were greater than the MPRAGE workflow. The wLOA was 28.7% for the comparison of the MPF and MPFreg workflows, compared to 36.9 for the comparison between the MPFreg and MPRAGE workflows, and 42% for the comparison between the MPF and MPRAGE workflows. This pattern was different in subcortical regions, where the MPF and MPFreg had a highly consistent agreement, with minimal bias and a wLOA of 15.6%. In subcortical regions, the MPRAGE workflow yielded volumes that were, on average, larger than those produced by the other two workflows. The wLOA was 39.4% for the comparison between the MPF and MPRAGE workflows and 39.8% for comparison between the MPFreg and MPRAGE workflows.

**Figure 9.**
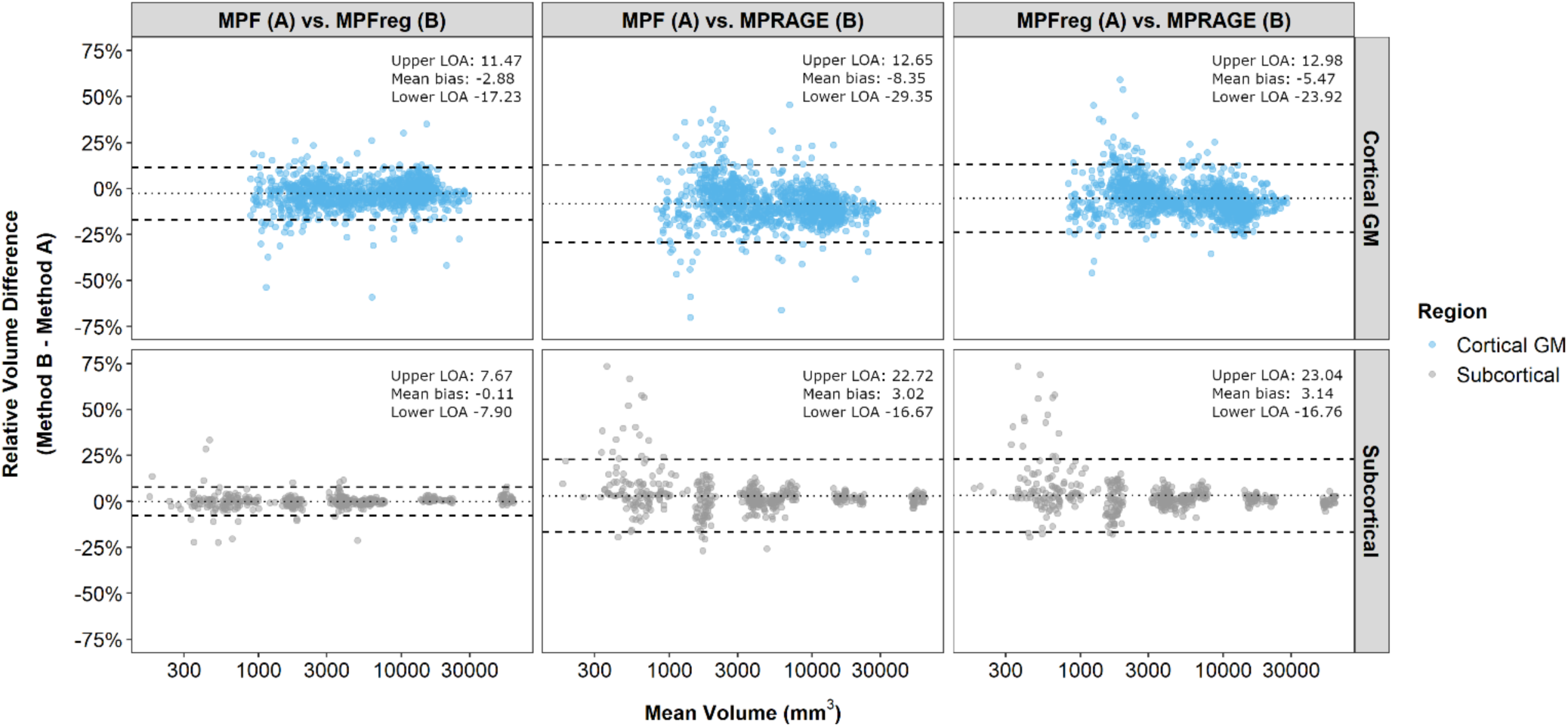
Bland-Altman plots for relative difference in volume for comparisons between workflows for cortical GM and subcortical regions. Dotted and dashed lines indicate mean bias and 95% limits of agreement (LOA), respectively.

### Scan-rescan repeatability of MPF measurements for different processing workflows

Within subject test-retest coefficient of variation (CVw) for mean regional MPF values are shown in Figure 10. In the cerebrum and lobe GM and WM regions, CVw for mean MPF did not differ significantly between workflows. CVws for the cerebrum ranged from 1.3%-1.7% in GM and were equal to 1.3% in WM. Of the lobes, the occipital lobe had the largest CVws, ranging from 2.8%-3.3% in GM and 2.6%-3.4% in WM. For the other three lobes, CVw ranged from 1.8%-2.9% in GM and from 1.6%-2.1% in WM. Significant differences between workflows were observed in a relatively small number of GM and WM cortical parcels. For all workflows, CVw for cortical parcels in GM and WM ranged from 2-7%. For subcortical regions, CVw values ranged from 2-6%, and significant differences between workflows were found for 6 out of the 16 subcortical regions. The regions with consistently larger CVw were the temporal pole, entorhinal, and pars orbitalis parcels, and the brainstem, hippocampus, amygdala, and nucleus accumbens.

**Figure 10.**
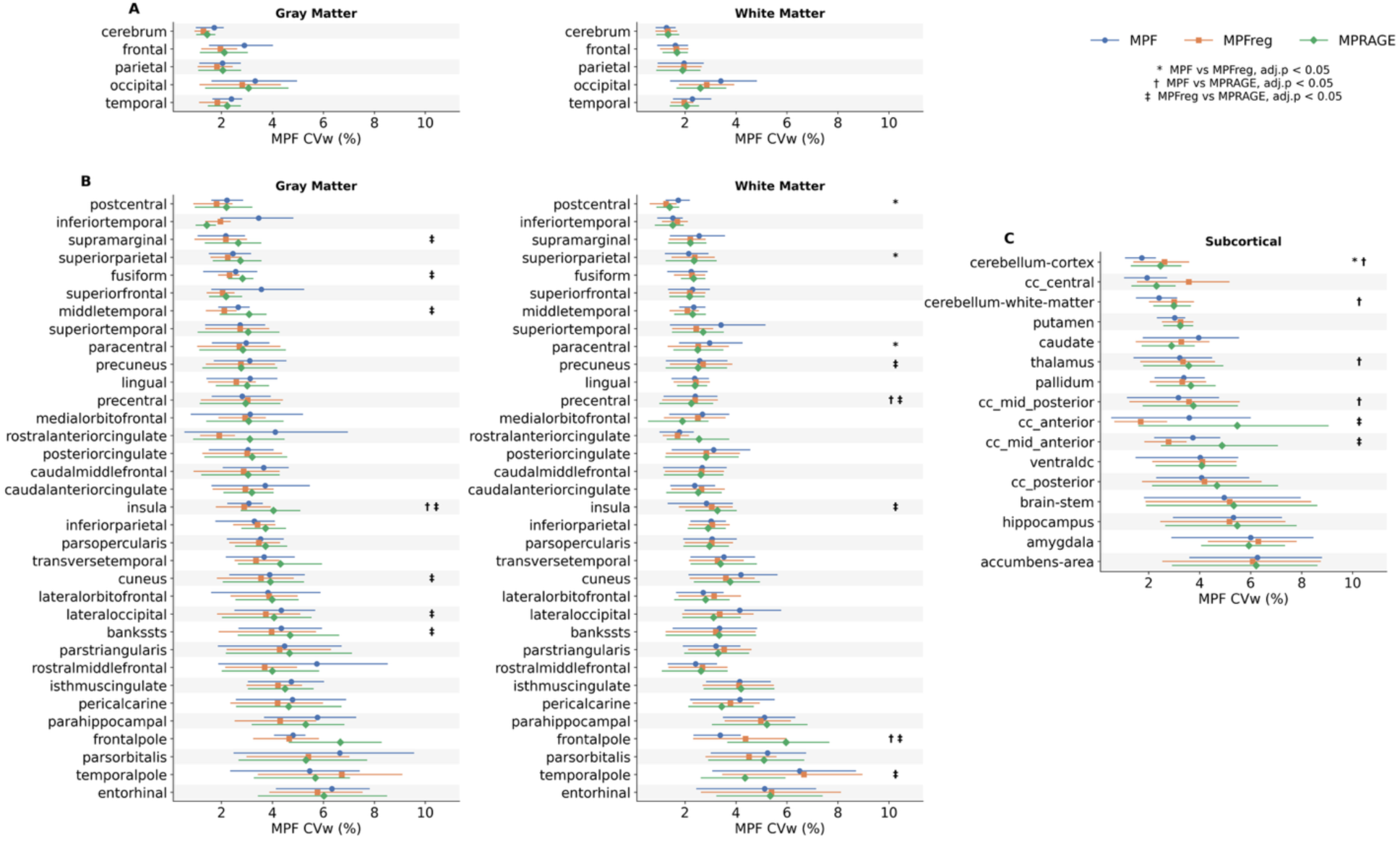
Within-subject test-retest coefficient of variation (CVw, %) for mean MPF values for the three processing workflows, shown for cerebrum and lobe GM and WM (A), cortical parcel GM and WM (B), and subcortical regions (C). Color and marker shape indicate workflow (MPF: blue circle; MPFreg: orange square; MPRAGE: green diamond). Horizontal lines represent confidence intervals. Symbols to the right indicate the presence of statistical significance in mean MPF values between workflows.

Within subject test-retest coefficient of variation (CVw) for regional volumes are shown in Figure 11. For the cerebrum and lobes, CVw ranged from 0.5-1.5% for the MPRAGE workflow, compared to 1.5-3.6% for the MPF workflow and 2.0-2.8% for the MPFreg workflow. The MPRAGE workflow had significantly lower CVw than the MPF, MPFreg, or both workflows in the cerebrum, all lobes except the occipital lobe, and most GM parcels. CVw for the MPRAGE workflow did not exceed 5% for any GM parcels, whereas it ranged up to 10% for the MPF workflow and up to 9% for the MPFreg workflow. Significant differences between workflows were observed in 4 of 16 subcortical regions, and 4 subcortical regions had CVw exceeding 5% for at least one workflow. Mean CVws were large and associated confidence intervals were comparatively wide as compared to other regions for the MPFreg or MPRAGE workflows in the central, mid-anterior, anterior, and mid-posterior segments of the corpus callosum.

**Figure 11.**
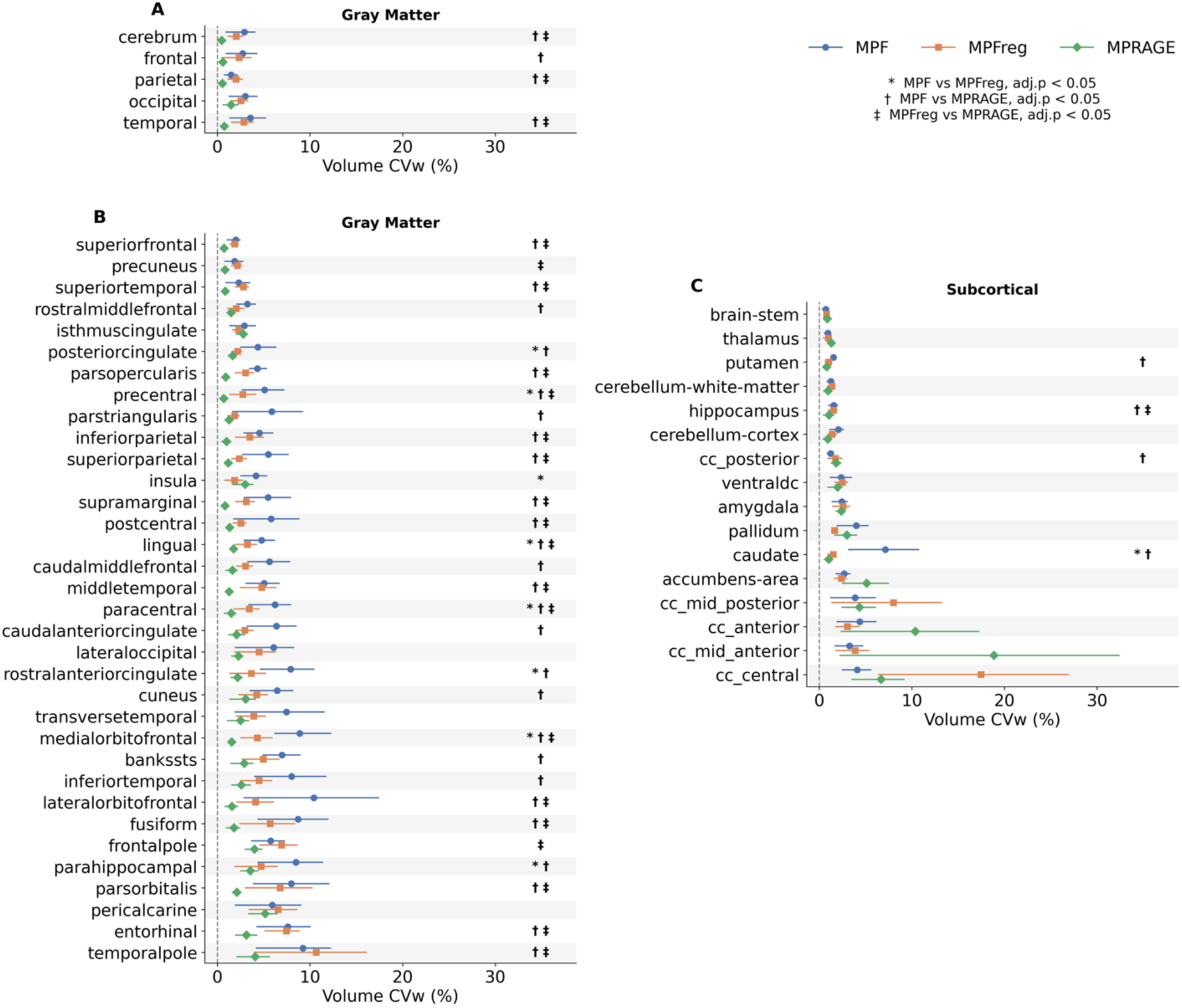
Within-subject test-retest coefficient of variation (CVw, %) for volume estimates for the three processing workflows, shown for cerebrum and lobe GM regions (A), cortical parcels (B), and subcortical regions (C). Color and marker shape indicate workflow (MPF: blue circle; MPFreg: orange square; MPRAGE: green diamond). Horizontal lines represent confidence intervals. Dashed vertical line indicates CVw = 0. Symbols to the right indicate the presence of statistical significance between processing workflows.

## Discussion

Overall, the three workflows produced generally comparable regional MPF and volume estimates, but the estimates were not interchangeable. Reliability of estimates of mean MPF was higher in WM and subcortical regions, where it was generally excellent, than in cortical GM, where it was more regionally dependent and ranged from poor to excellent. In both GM and WM, reliability for mean MPF was lowest in regions near air-tissue boundaries. For volume estimates, reliability in cortical parcels was generally good to excellent for comparisons between the MPF and MPFreg workflows, and between the MPFreg and MPRAGE. In contrast, the MPF workflow showed moderate to poor reliability for volume estimates in a larger number of cortical parcels. Just as observed for mean MPF, reliability across methods for volume estimates was lowest in regions near air-tissue boundaries. In subcortical regions, reliability was highest in comparisons between the MPFreg and MPF workflows, and between the MPFreg and MPRAGE workflows. In comparison, reliability was lower in numerous regions for the comparison between the MPF and MPRAGE workflow.

For estimation of mean MPF, systematic biases between workflows were observed in WM and subcortical regions. In WM, the MPF and MPFreg workflows yielded consistently larger values than the MPRAGE workflow, and the MPF workflow yielded consistently larger values than the MPFreg workflow. In subcortical regions, the same pattern was observed for most regions. In GM, no bias was observed between the MPF and MPFreg workflows, while both workflows resulted in predominately larger values than the MPRAGE workflow. For regional volumes, a systematic bias between workflows was found for the cerebrum WM region, with the MPRAGE workflow yielding larger values than the MPFreg and MPF workflows, and the MPFreg workflow yielding larger values than the MPF workflow. In GM, the MPF and MPFreg workflows yielded larger values than the MPRAGE workflow in most regions, and the MPFreg workflow yielded larger values than the MPF workflow in many regions. In subcortical regions, the MPRAGE workflow yielded larger values than the MPF and MPFreg workflows in most regions, while differences between the MPF and MPFreg workflows were smaller and inconsistent in direction.

To the best of our knowledge, this is the first report on scan-rescan repeatability of regional MPF measurements based on automated brain segmentation and parcellation. Two earlier studies (Smirnova et al., 2021; Yarnykh et al., 2020) examined repeatability of whole-brain WM and GM MPF values measured using a Markov random field segmentation algorithm (Zhang et al., 2001). These studies reported CVw values on the order of 1-2%, which is consistent with the presented results (∼1.5%). The lobe-level MPF measurements we report also appear highly reproducible, with CVw of approximately 2-3%. As expected, variability for the aggregate regions (cerebrum or lobes) was often lower than that for cortical parcels and subcortical regions. At the same time, the majority of cortical GM and WM as well as subcortical structures showed reasonably good repeatability with CVw less than 5%. Repeatability findings were not affected by tissue type, as they were generally similar between WM and GM in both cortical and subcortical regions. For mean MPF, very little difference in repeatability was observed between workflows. The major factor affecting regional MPF repeatability was the anatomic region itself. Reduced repeatability (larger CVw) was mainly observed in regions prone to magnetic susceptibility effect due to proximity to air-tissue interfaces, such as the temporal and frontal poles, the entorhinal cortex, and the amygdala. Similar anatomical patterns of increased variability for volumetric and surface measures derived from FreeSurfer have been previously reported (Knussmann et al., 2022). Higher variability was also observed in very small anatomic structures, such as the nucleus accumbens.

For volume measurements, the MPRAGE workflow consistently had the smallest CVw (generally <5% for parcels and subcortical regions, and <1% at the whole-brain and lobe levels), which is generally similar to previous literature on the repeatability of FreeSurfer parcellation (Jovicich et al., 2009; Knussmann et al., 2022; Okubo et al., 2016; Seiger et al., 2015; Wonderlick et al., 2009; Yan et al., 2020). Variability was significantly higher for the MPF and MPFreg workflows than for the MPRAGE workflow in most regions. Larger uncertainties in the estimation of MPF-based volumes may be associated with broadening of tissue boundaries caused by sub-voxel displacements of the source images used to compute MPF maps. Such displacements are unavoidable for synthetic imaging techniques that utilize multiple sequentially acquired source images, as these displacements cannot usually be completely corrected by registration algorithms. This observation also can be a manifestation of the slightly lower spatial resolution of the MPF maps as compared to the MPRAGE images (1.25 mm^3^ vs 1 mm^3^) used in this study. One exception was the corpus callosum segments, where the MPFreg or MPRAGE workflows showed poor repeatability, reflected by relatively large CVw values (up to 18%). It is also noticeable that MPF-based corpus callosum segmentation resulted in less variable mean MPF estimates as compared to the MPRAGE-based ones, and that the corpus callosum segments were the regions with the largest disagreement between MPF-based and MPRAGE-based workflows in terms of both mean MPF and volume measurements. While we cannot offer a clear explanation of this observation, practically it suggests that MPF could be a preferable modality for the studies specifically focused on measurements for corpus callosum segments.

Consistency ICC values for mean MPF were generally lower for GM than WM, but only for comparisons involving MPRAGE; MPF and MPFreg showed comparable ICC values across both tissue types. Bland-Altman analysis revealed wider wLOA for GM than WM in the majority of regions for all three comparisons, indicating greater workflow disagreement in GM regardless of which workflows were compared. However, this wider LOA only translated into lower ICC when MPRAGE images were involved in the workflow, suggesting that the disagreement between MPRAGE and MPF-based measures in GM is too large for the workflows to be used interchangeably, whereas the disagreement between MPF and MPFreg remains acceptable relative to between-subject biological variability. This likely reflects lower absolute MPF values in GM, where small processing workflow differences represent a proportionally larger error, and greater susceptibility to partial volume effects at the GM/CSF and GM/WM boundaries. This is most relevant to studies relying on individual-level estimates, such as brain-behavior correlations, predictive models, longitudinal within-subject analyses, and classification approaches, where noise in individual measurements reduces sensitivity and stability. The findings of high repeatability in this study suggest that variations in reliability across tissue types and workflows is not attributable to measurement instability. The variability in reliability and bias across regions for most workflow comparison also indicate that any study should utilize a single, consistent processing workflow for all data.

This study has several limitations. First, the sample was small. Second, the sample was comprised of adults. Differences between processing workflows may be more pronounced in pediatric or clinical populations where increased head motion and reduced compliance are more common. Motion-related artifacts can degrade image quality and introduce spatial inconsistencies that propagate through processing steps. As a result, both systematic bias and differences in relative values across participants between workflows may be amplified in these populations relative the adult sample of the present study.

In conclusion, this study demonstrated that whole-brain 3D MPF maps can be used as input images for FreeSurfer parcellation and enable high repeatability of mean MPF estimates across the majority of WM, GM, and subcortical structures. Further improvement in reliability of MPF measurements, particularly in cortical GM, can be achieved by applying rigid registration of source images prior to MPF map reconstruction and measuring MPF using regional masks derived from registered high-resolution structural T1-weighted images rather than native MPF maps. Application of MPF maps instead of T1-weighted images for the purpose of estimating brain volume should be done with caution. For both types of measurements, workflow choice may introduce small but significant biases in regional, lobar, and whole-brain MPF values. This should be taken into account for studies that involve comparisons between estimates at the individual subject level, such as studies focused on clinical and behavioral correlates of brain myelination.

## Data and Code Availability

The data for this study and the code used in the analyses are available upon request.

## Author Contributions

Conceptualization: V.Y Methodology: D.H., V.Y. Investigation: D.H., N.M.C. Visualization: N.M.C. Supervision: V.Y. Writing original draft: N.M.C. Writing—review & editing: N.M.C., D.H., V.Y.

## Funding

This study was supported by the National Institutes of Health grant R01NS136227. Development of software for MPF map reconstruction was supported by National Institutes of Health grant R24NS104098.

## Declaration of Competing Interest

The authors declare that there is no conflict of interest regarding the publication of this article.

## Notes

### Competing Interest Statement

The authors have declared no competing interest.

## References

Avants, B. B., Tustison, N. J., Song, G., Cook, P. A., Klein, A., & Gee, J. C. (2011). A reproducible evaluation of ANTs similarity metric performance in brain image registration. NeuroImage, 54(3), 2033–2044. 10.1016/j.neuroimage.2010.09.025

Benjamini, Y., & Hochberg, Y. (1995). Controlling the false discovery rate: A practical and powerful approach to multiple testing. Journal of the Royal Statistical Society: Series B (Methodological), 57(1), 289–300. 10.1111/j.2517-6161.1995.tb02031.x

Bland, J. M., & Altman, D. G. (1986). Statistical methods for assessing agreement between two methods of clinical measurement. The Lancet, 1(8476), 307–310.

Chubb, H., Karim, R., Roujol, S., Nuñez-Garcia, M., Williams, S. E., Whitaker, J., Harrison, J., Butakoff, C., Camara, O., Chiribiri, A., Schaeffter, T., Wright, M., O’Neill, M., & Razavi, R. (2018). The reproducibility of late gadolinium enhancement cardiovascular magnetic resonance imaging of post-ablation atrial scar: A cross-over study. Journal of Cardiovascular Magnetic Resonance, 20, 21. 10.1186/s12968-018-0438-y

Corrigan, N. M., Yarnykh, V. L., Hippe, D. S., Owen, J. P., Huber, E., Zhao, T. C., & Kuhl, P. K. (2021). Myelin development in cerebral gray and white matter during adolescence and late childhood. NeuroImage, 227. 10.1016/j.neuroimage.2020.117678

Corrigan, N. M., Yarnykh, V. L., Huber, E., Zhao, T. C., & Kuhl, P. K. (2022). Brain myelination at 7 months of age predicts later language development. NeuroImage, 263. 10.1016/j.neuroimage.2022.119641

Desikan, R. S., Ségonne, F., Fischl, B., Quinn, B. T., Dickerson, B. C., Blacker, D., Buckner, R. L., Dale, A. M., Maguire, R. P., Hyman, B. T., Albert, M. S., & Killiany, R. J. (2006). An automated labeling system for subdividing the human cerebral cortex on MRI scans into gyral based regions of interest. NeuroImage, 31(3), 968–980. 10.1016/j.neuroimage.2006.01.021

Filimonova, E. A., Pashkov, A. A., Yarnykh, V. L., Schukina, M. I., Zaitsev, B. A., Martirosyan, A. V., Moysak, G. I., & Rzaev, J. A. (2025). Assessment of Trigeminal Nerve Root Demyelination in Patients with Primary Trigeminal Neuralgia Using Macromolecular Proton Fraction Imaging. American Journal of Neuroradiology, 46(3), 602–610. 10.3174/ajnr.A8545

Fischl, B. (2012). FreeSurfer. NeuroImage, 62(2), 774–781. 10.1016/j.neuroimage.2012.01.021

FreeSurfer. (2009). FreeSurfer Suggested Morphometry Protocols. https://surfer.nmr.mgh.harvard.edu/fswiki/FreeSurferWiki?action=AttachFile&do=get&target=FreeSurfer_Suggested_Morphometry_Protocols.pdf

Gamer, M., Lemon, J., Fellows, I., & Singh, P. (2026). irr: Various Coefficients of Interrater Reliability and Agreement (Version 0.85) [Computer software]. https://cran.r-project.org/web/packages/irr/index.html

Gusakova, S., Smirnova, L., Borodin, O., Epimakhova, E., Seregin, A., & Yarnykh, V. (2026). Macromolecular Proton Fraction Reveals Divergent White Matter Myelination in Bipolar Disorder and Unipolar Recurrent Depression. Bioengineering, 13(1), 78. 10.3390/bioengineering13010078

Helms, G., & Dechent, P. (2009). Increased SNR and reduced distortions by averaging multiple gradient echo signals in 3D FLASH imaging of the human brain at 3T. Journal of Magnetic Resonance Imaging: JMRI, 29(1), 198–204. 10.1002/jmri.21629

Huang, F. L. (2018). Using Cluster Bootstrapping to Analyze Nested Data With a Few Clusters. Educational and Psychological Measurement, 78(2), 297–318. 10.1177/0013164416678980

Hyslop, N. P., & White, W. H. (2009). Estimating Precision Using Duplicate Measurements. Journal of the Air & Waste Management Association, 59(9), 1032–1039. 10.3155/1047-3289.59.9.1032

Jovicich, J., Czanner, S., Han, X., Salat, D., van der Kouwe, A., Quinn, B., Pacheco, J., Albert, M., Killiany, R., Blacker, D., Maguire, P., Rosas, D., Makris, N., Gollub, R., Dale, A., Dickerson, B. C., & Fischl, B. (2009). MRI-derived measurements of human subcortical, ventricular and intracranial brain volumes: Reliability effects of scan sessions, acquisition sequences, data analyses, scanner upgrade, scanner vendors and field strengths. NeuroImage, 46(1), 177–192. 10.1016/j.neuroimage.2009.02.010

Khodanovich, M., Kamaeva, D., Usova, A., Pashkevich, V., Moshkina, M., Obukhovskaya, V., Kataeva, N., Levina, A., Tumentceva, Y., Shadrina, M., Ranzaeva, A., Vasilieva, S., Schastnyy, E., Naumova, A., & Svetlik, M. (2025). Demyelination and Cognitive Performance in Long COVID Patients with Insomnia and/or Depression. International Journal of Molecular Sciences, 26(24). 10.3390/ijms262412141

Khodanovich, M., Kisel, A. A., Akulov, A. E., Atochin, D. N., Kudabaeva, M. S., Glazacheva, V. Y., Svetlik, M. V., Medvednikova, Y. A., Mustafina, L. R., & Yarnykh, V. L. (2018). Quantitative assessment of demyelination in ischemic stroke in vivo using macromolecular proton fraction mapping. Journal of Cerebral Blood Flow and Metabolism, 38(5), 919–931. 10.1177/0271678X18755203

Kisel, A. A., Naumova, A. V., & Yarnykh, V. L. (2022). Macromolecular Proton Fraction as a Myelin Biomarker: Principles, Validation, and Applications. Frontiers in Neuroscience, 16, 819912. 10.3389/fnins.2022.819912

Knussmann, G. N., Anderson, J. S., Prigge, M. B. D., Dean, D. C., Lange, N., Bigler, E. D., Alexander, A. L., Lainhart, J. E., Zielinski, B. A., & King, J. B. (2022). Test-retest reliability of FreeSurfer-derived volume, area and cortical thickness from MPRAGE and MP2RAGE brain MRI images. Neuroimage: Reports, 2(2). 10.1016/j.ynirp.2022.100086

Koo, T. K., & Li, M. Y. (2016). A Guideline of Selecting and Reporting Intraclass Correlation Coefficients for Reliability Research. Journal of Chiropractic Medicine, 15(2), 155–163. 10.1016/j.jcm.2016.02.012

Korostyshevskaya, A. M., Prihod’ko, I. Y., Savelov, A. A., & Yarnykh, V. L. (2019). Direct comparison between apparent diffusion coefficient and macromolecular proton fraction as quantitative biomarkers of the human fetal brain maturation. Journal of Magnetic Resonance Imaging, 50(1), 52–61. 10.1002/jmri.26635

Korostyshevskaya, A. M., Savelov, A. A., Papusha, L. I., Druy, A. E., & Yarnykh, V. L. (2018). Congenital medulloblastoma: Fetal and postnatal longitudinal observation with quantitative MRI. Clinical Imaging, 52, 172–176. 10.1016/j.clinimag.2018.06.001

Okubo, G., Okada, T., Yamamoto, A., Kanagaki, M., Fushimi, Y., Okada, T., Murata, K., & Togashi, K. (2016). MP2RAGE for deep gray matter measurement of the brain: A comparative study with MPRAGE. Journal of Magnetic Resonance Imaging, 43(1), 55–62. 10.1002/jmri.24960

Seiger, R., Hahn, A., Hummer, A., Kranz, G. S., Ganger, S., Küblböck, M., Kraus, C., Sladky, R., Kasper, S., Windischberger, C., & Lanzenberger, R. (2015). Voxel-based morphometry at ultra-high fields. A comparison of 7T and 3T MRI data. NeuroImage, 113, 207–216. 10.1016/j.neuroimage.2015.03.019

Smirnova, L. P., Yarnykh, V. L., Parshukova, D. A., Kornetova, E. G., Semke, A. V., Usova, A. V., Pishchelko, A. O., Khodanovich, M. Y., & Ivanova, S. A. (2021). Global hypomyelination of the brain white and gray matter in schizophrenia: Quantitative imaging using macromolecular proton fraction. Translational Psychiatry, 11(1). 10.1038/s41398-021-01475-8

Smith, S. M., Jenkinson, M., Woolrich, M. W., Beckmann, C. F., Behrens, T. E. J., Johansen-Berg, H., Bannister, P. R., De Luca, M., Drobnjak, I., Flitney, D. E., Niazy, R. K., Saunders, J., Vickers, J., Zhang, Y., De Stefano, N., Brady, J. M., & Matthews, P. M. (2004). Advances in functional and structural MR image analysis and implementation as FSL. *NeuroImage*, Mathematics in Brain Imaging, 23, S208– S219. 10.1016/j.neuroimage.2004.07.051

Sui, Y. V., McKenna, F., Bertisch, H., Storey, P., Anthopolos, R., Goff, D. C., Samsonov, A., & Lazar, M. (2022). Decreased basal ganglia and thalamic iron in early psychotic spectrum disorders are associated with increased psychotic and schizotypal symptoms. Molecular Psychiatry, 27(12), 5144–5153. 10.1038/s41380-022-01740-2

van Houdt, P. J., Li, S., Yang, Y., & van der Heide, U. A. (2024). Quantitative MRI on MR-Linacs: Towards Biological Image-Guided Adaptive Radiotherapy. Seminars in Radiation Oncology, 34(1), 107–119. 10.1016/j.semradonc.2023.10.010

Wonderlick, J. S., Ziegler, D. A., Hosseini-Varnamkhasti, P., Locascio, J. J., Bakkour, A., van der Kouwe, A., Triantafyllou, C., Corkin, S., & Dickerson, B. C. (2009). Reliability of MRI-derived cortical and subcortical morphometric measures: Effects of pulse sequence, voxel geometry, and parallel imaging. NeuroImage, 44(4), 1324–1333. 10.1016/j.neuroimage.2008.10.037

Yan, S., Qian, T., Maréchal, B., Kober, T., Zhang, X., Zhu, J., Lei, J., Li, M., & Jin, Z. (2020). Test-retest variability of brain morphometry analysis: An investigation of sequence and coil effects. Annals of Translational Medicine, 8(1), 12–12. 10.21037/atm.2019.11.149

Yarnykh, V. L. (2010). Optimal radiofrequency and gradient spoiling for improved accuracy of T1 and B1 measurements using fast steady-state techniques. Magnetic Resonance in Medicine, 63(6), 1610– 1626. 10.1002/mrm.22394

Yarnykh, V. L. (2012). Fast macromolecular proton fraction mapping from a single off-resonance magnetization transfer measurement. Magnetic Resonance in Medicine, 68(1), 166–178. 10.1002/mrm.23224

Yarnykh, V. L. (2016). Time-efficient, high-resolution, whole brain three-dimensional macromolecular proton fraction mapping. Magnetic Resonance in Medicine, 75(5), 2100–2106. 10.1002/mrm.25811

Yarnykh, V. L. (2021). Data-Driven Retrospective Correction of B1Field Inhomogeneity in Fast Macromolecular Proton Fraction and R1Mapping. IEEE Transactions on Medical Imaging, 40(12), 3473–3484. 10.1109/TMI.2021.3088258

Yarnykh, V. L., Bowen, J. D., Samsonov, A., Repovic, P., Mayadev, A., Qian, P., Gangadharan, B., Keogh, B. P., Maravilla, K. R., & Henson, L. K. J. (2015). Fast whole-brain three-dimensional macromolecular proton fraction mapping in multiple sclerosis. Radiology, 274(1), 210–220. 10.1148/radiol.14140528

Yarnykh, V. L., Kisel, A. A., & Khodanovich, M. Y. (2020). Scan–Rescan Repeatability and Impact of B0 and B1 Field Nonuniformity Corrections in Single-Point Whole-Brain Macromolecular Proton Fraction Mapping. Journal of Magnetic Resonance Imaging, 51(6), 1789–1798. 10.1002/jmri.26998

Yarnykh, V. L., Prihod’ko, I. Y., Savelov, A. A., & Korostyshevskaya, A. M. (2018). Quantitative assessment of normal fetal brain myelination using fast macromolecular proton fraction mapping. American Journal of Neuroradiology, 39(7). 10.3174/ajnr.A5668

Zhang, Y., Brady, M., & Smith, S. (2001). Segmentation of brain MR images through a hidden Markov random field model and the expectation-maximization algorithm. IEEE Transactions on Medical Imaging, 20(1), 45–57. 10.1109/42.906424

Zhao, T. C., Corrigan, N. M., Yarnykh, V. L., & Kuhl, P. K. (2022). Development of executive function-relevant skills is related to both neural structure and function in infants. Developmental Science, 25(6), e13323. 10.1111/desc.13323

